# Quantitative examination of five stochastic cell-cycle and cell-size control models for *Escherichia coli* and *Bacillus subtilis*

**DOI:** 10.1101/2021.06.06.447266

**Authors:** Guillaume Le Treut, Fangwei Si, Dongyang Li, Suckjoon Jun

## Abstract

We examine five quantitative models of the cell-cycle and cell-size control in *Escherichia coli* and *Bacillus subtilis* that have been proposed over the last decade to explain single-cell experimental data generated with high-throughput methods. After presenting the statistical properties of these models, we test their predictions against experimental data. Based on simple calculations of the defining correlations in each model, we first dismiss the stochastic Helmstetter-Cooper model and the Initiation Adder model, and show that both the Replication Double Adder and the Independent Double Adder model are more consistent with the data than the other models. We then apply a recently proposed statistical analysis method and obtain that the Independent Double Adder model is the most likely model of the cell cycle. By showing that the Replication Double Adder model is fundamentally inconsistent with size convergence by the adder principle, we conclude that the Independent Double Adder model is most consistent with the data and the biology of bacterial cell-cycle and cell-size control. Mechanistically, the Independent Adder Model is equivalent to two biological principles: (i) balanced biosynthesis of the cell-cycle proteins, and (ii) their accumulation to a respective threshold number to trigger initiation and division.

## 1. Introduction

Quantitative microbial physiology is marked by close interactions between experiment and modeling since its birth in the mid 20th century (see (1) for a review of the history with extensive literature). In particular, bacterial cell-size and cell-cycle control has enjoyed rejuvenated interests in modeling with the advent of microfluidics techniques that allow tracking of thousands of individual cells over a hundred division cycles (see, for example, (2–5). Re-emerged from the new single-cell data is the adder principle (6–8), which states that individual cells grow by adding a fixed size from birth to division, independently from their size at birth. This principle has characteristic repercussions on cell size homeostasis. Specifically, upon perturbation, the cell size at birth relaxes toward its steady-state value according to a first-order recurrence relation with a correlation coefficient equal to 1/2 (8–10).

Although the adder principle was originally proposed and statistically tested almost three decades ago by Koppes and colleagues before its recent revival (9), its mechanistic origin has remained elusive until recently because direct experimental tests were not available for a long time (1, 11). Several models have been proposed so far (6, 8, 11–20), and we expect a consensus to emerge as more experimental data become available.

The main purpose of this article is to derive and present steady-state statistical properties of quantitative bacterial cell-cycle and cell-size control models that we are currently aware of and, where relevant, critically examine them against single-cell data from our lab’s mother machine experiments accumulated over the last decade in *E. coli* and *B. subtilis*. These models are (i). the stochastic Helmstetter-Cooper model (sHC) (11), (ii). the initiation adder (IA) model (10, 13, 21), (iii). the replication double adder (RDA) model (17), (iv). the independent double adder (IDA) model (11), and (v). the concurrent cell-cycle processes (CCCP) model (22) and its stochastic version (14).

Some of these models are graphically illustrated in Figure 1. Briefly, the sHC model is a literal extension of the textbook Helmstetter Cooper model by allowing independent Gaussian fluctuations to each of the initiation mass, the *τ*_cyc_=C+D period (from initiation to division), and the cell elongation rate. The IA model assumes that replication initiation is the sole implementation point of cell-size control, and division is strictly coupled to initiation such that division is triggered after fixed *τ*_cyc_=C+D minutes have elapsed since initiation. The RDA model is similar to the IA model in that it also assumes that initiation is the reference point for cell-size control. Its main difference from the IA model is that it assumes division is triggered after the cell elongates a constant length per origin of replication *δ*_id_, rather than a constant time, since initiation. In other words, *δ*_id_ is the added size during the C+D period. Both the IA and the RDA models assume the initiation adder, i.e., the cell growth by a nearly fixed size per replication origin between two consecutive initiation cycles, irrespective of the cell size at initiation (initiation mass).

**Figure 1:**
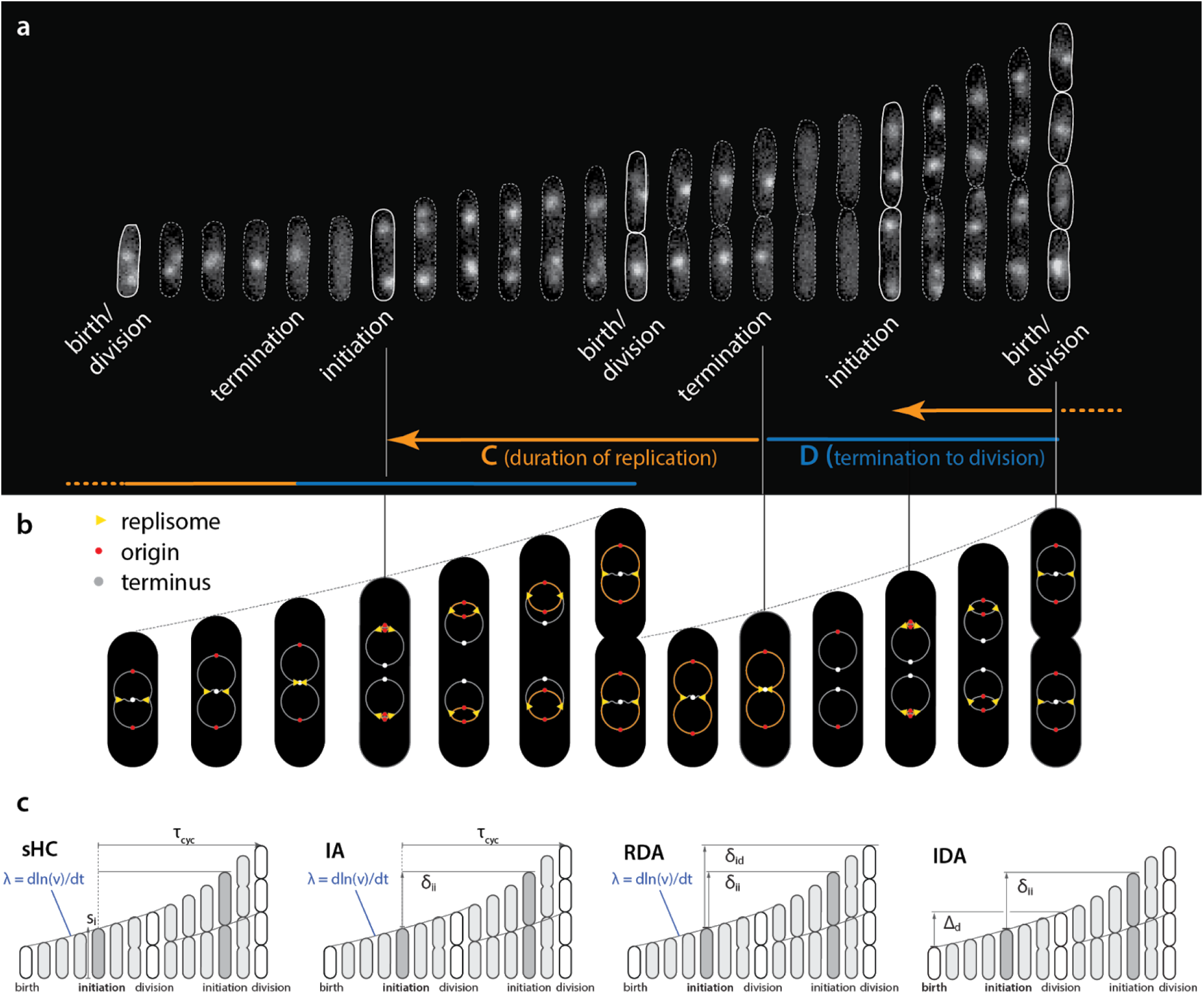
Physiological parameters that can be measured from single-cell experiments. **a**. Time-lapse images of a single *Escherichia coli* cell growing in a microfluidics channel. The cell boundaries are segmented from phase contrast images whereas the replication forks are visualized using a functional fluorescently labeled replisome protein (DnaN-YPet). **b**. Multifork replication: in most growth conditions, several replication cycles overlap. The direction of the arrows is not the direction of time, but to illustrate that the HC model’s core idea is to trace replication initiation backward in time by C+D from division. **c**. Four models of *E. coli* cell cycle and their control variables, which can be measured from single-cell experiments. The sHC model describes cell size and cell cycle using three parameters: elongation rate *λ* = = dln(*l*)/d*t*, where *l* is the cell length (not shown), *τ*_cyc_ = *C*+*D*, and the initiation size per origin of replication *s*_i_. The IA model uses *λ, τ*_cyc_ and the added size per origin of replication between consecutive replication initiation events *δ*_ii_. The RDA model uses *λ, δ*_ii_ and the added size per origin of replication from initiation to division *δ*_id_. The IDA model uses *λ, δ*_ii_ and the added size from birth to division *Δ*_d_. Note that both *δ*_id_ and *τ*_cyc_ can span multiple generations.

The IDA model also states that initiation and division are independently controlled by their own respective initiator proteins. However, the IDA model is based on mechanistic assumptions that these proteins are produced in a balanced manner (i.e., their production rate is the same as the growth rate of the cell), and initiation and division are triggered when the cell has accumulated the initiator proteins to their respective threshold numbers. The CCCP model states that replication cycle and division cycles progress independently, but checkpoints or their equivalent are activated to ensure cell division (22).

This article is structured as follows. In Section 2, we summarize the five models and derive some of their statistical properties. In Section 3, we test the predictions of these models against the data. In Section 4, we critically examine one of the recent correlation analysis methods (the *I*-value analysis) used to justify the RDA model. We conclude that the IDA model is as of today the model most consistent with data, which also provides a falsifiable mechanistic picture.

## 2. Statistical properties of five bacterial cell-size and cell-cycle control models

### 2.1 The stochastic Helmstetter-Cooper model (sHC)

The original HC model (23) is based on the experimental observation that the average duration of chromosome replication (“*C* period”) can be longer than the average doubling time of the cells in fast-growing *E. coli*. In such growth conditions, *E. coli* must initiate a new round of replication before the ongoing replication cycle is completed. The core of the HC model is the recipe to trace replication initiation backward by *τ*_cyc_=*C*+*D* > *τ* minutes starting from cell division during overlapping cell cycles (Figure 1).

Thus, the HC model introduces three control parameters for a complete description of replication and division cycles: two temporal parameters (the doubling time *τ* and the duration of cell cycle *τ*_cyc_ = *C*+*D*) and one spatial parameter (e.g., cell size at division or initiation). It was Donachie who showed that, if (i) *τ*_cyc_ = *C*+*D* is invariant under different nutrient conditions and (ii) the average cell size increases exponentially with respect to the nutrient-imposed growth rate λ = 2/ln*τ* as *S* = *s*_i_ exp(*α λ*) (where *s*_i_ and *α* are constant, and *S* is the average cell size of steady-state population), then the cell size at initiation per replication origin *s*_i_ (or, the “initiation mass”) must be mathematically invariant in all growth conditions (24). This result was later generalized to all steady-state growth conditions with and without growth inhibition (25).

Since the original HC model is deterministic and can be defined in terms of *λ, τ*_cyc_ and *s*_i_ (or division size *S*_d_), one possible extension to a stochastic version is by making the three physiological variables stochastic. Together, they completely determine cell sizes including the size at division, assuming perfectly symmetric division (Figure 1). For simplicity, we draw *λ, τ*_cyc_ and *s*_i_ at cell birth from a multivariate Gaussian distribution, which also encodes cross- and mother-daughter correlations in the covariances matrix (11).

The recursion relation for the cell size at division in this “stochastic” Helmstetter-Cooper (sHC) model can be written as follows:

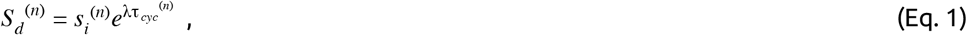

where *n* denotes the generation index. If we assume that cells elongate exponentially at the growth rate *λ* (2), *the number of overlapping cell cycles p*+1 is completely determined by the relation:

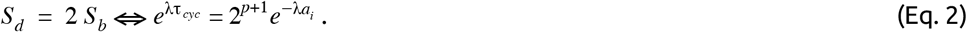

where *a*_i_ is the time duration elapsed between cell birth and replication initiation. It follows that *p* is the integer part of *τ*_cyc_/*τ*, where *τ* = ln2 / *λ* is the generation time, so that *p*+1 is the number of overlapping cell cycles (unless noted otherwise we will adopt the convention that *X*^(n)^ denotes the value of a physiological variable in generation *n* whereas *X* is the average over the whole lineage).

In the sHC model, consecutive sizes at initiation are correlated through *ρ*_i_ = *ρ*(*s*_i_^(n)^, *s*_i_^(n+1)^) = 0, where *ρ*(*A,B*) stands for the Pearson correlation coefficient between variables *A* and *B*. In the absence of mother-daughter correlations for all three physiological variables, the cell should behave as a sizer, *ρ*_d_ = *ρ*(*S*_d_^(n)^, *S*_d_^(n+1)^) = 0. However, additional cross- or auto-correlations among *λ, τ*_cyc_ and *s*_i_ (such as cross-correlations between *s*_i_^(n)^ and *τ*_cyc_^(n)^ and/or mother-daughter correlations between *s*_i_^(n)^ and *s*_i_^(n+1)^) can have a non-trivial effect on size homeostasis. Analytical expressions for *ρ*_i_, *ρ*_d_ and *ρ*_id_=(*s*_i_^(n)^, *s*_i_^(n+1)^) are derived in Appendix C. Importantly, *ρ*_d_ is particularly sensitive to the mother-daughter initiation-size correlation *ρ*_i_ = *ρ*(*s*_i_^(n)^, *s*_i_^(n+1)^) in the sHC model (11). This observation motivated an experimental study aiming at perturbing *ρ*_i_ by periodic expression of DnaA to break the adder phenotype in *E. coli*. This prediction was refuted in the experiments (11), since despite the perturbations to *ρ*_i_ by periodic oscillations of *dnaA* expression level, *E. coli* maintained its size homeostasis following the adder behavior (11). An important conclusion from the oscillation experiments is that replication initiation and cell division are independently controlled in steady-state conditions in both *E. coli* and *Bacillus subtilis*, thus firmly refuting the particular version of the sHC model.

### 2.2. The initiation adder model (IA)

The IA model is a variant of the sHC model in which the constraint on the initiation mass (*s*_i_ = constant in all growth conditions) is replaced by an adder mechanism running between consecutive replication initiations (21, 26). Specifically, the cell initiates replication following the adder principle, i.e., the size added per origin between two consecutive initiation cycles, *δ*_ii_, is independent of the cell size at initiation (13). Yet, as in the sHC model, the IA model assumes that division is triggered after a fixed duration of time, *τ*_cyc_, has elapsed since initiation. The three stochastic control parameters in the IA model are therefore: *λ, τ*_cyc_ and *δ*_ii_. A given cell size at replication initiation determines the next replication initiation event and one division event.

The recursion relation for cell size at division is the same as in the sHC model (Eq. 1). However, this relation is complemented with the following adder recursion relation determining the cell size per origin at replication initiation:

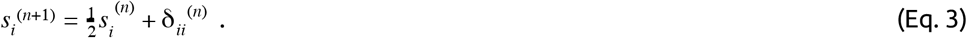

As before, *λ*^(n)^, *τ*_*cyc*_ ^(n)^ and *δ*_*ii*_ ^(n)^ are random variables associated with the *n*-th generation. To derive statistical properties of the IA model (Table 1), we will assume that they are independent Gaussian variables. At steady-state, Eq. 3 implies that *s*_i_ = 2*δ*_ii_. Therefore, the number of overlapping cell-cycles is also determined by Eq. 2 in the IA model (namely, *p*+1).

**Table 1.**
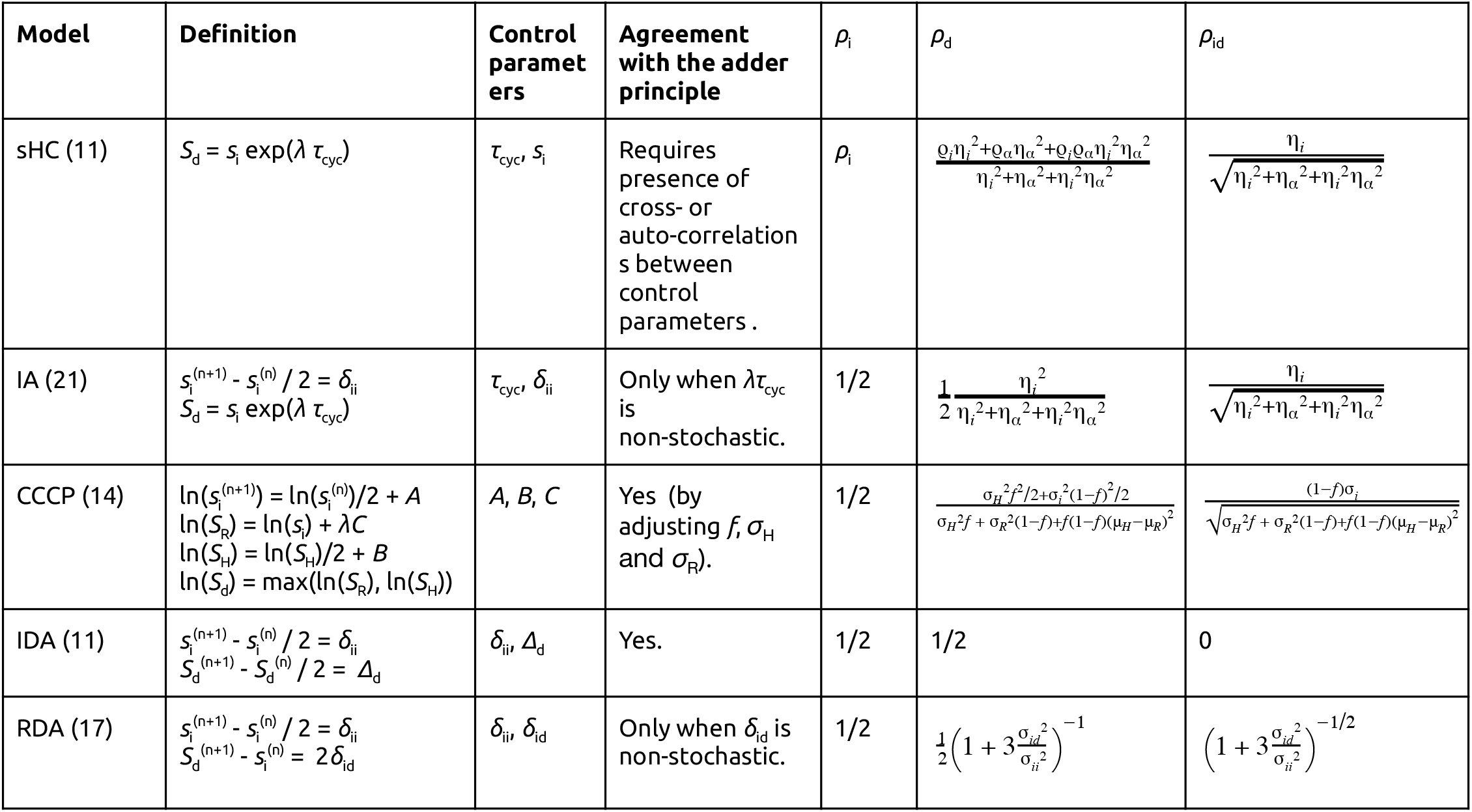
Summary of a few models of the *E. coli* cell cycle. The definition column indicates the equations defining the division and replication cycles. The control parameters are summarized in the next column. In the three rightmost columns we give the three correlations *ρ*_i_, *ρ*_d_ and *ρ*_id_. We have used the following variables: (i) *σ*_ii_^2^: variance of *δ*_ii_, (ii) *σ*_id_^2^: variance of *δ*_id_, (iii) *μ*_i_: mean of *s*_i_, (iv) *σ*_i_^2^: variance of *s*_i_, (v) *μ*_ɑ_: mean of *ɑ*=exp(*λτ*_cyc_), (vi) *σ*_ɑ_^2^: variance of *ɑ*, (vii) *η*_i_=*σ*_i_/*μ*_i_ is the coefficient of variation (CV) of *s*_i_, (viii) *η*_ɑ_=*σ*_ɑ_/*μ*_ɑ_ is the CV of *ɑ*, (ix) *μ*_H_: mean of ln(*S*_H_), (x) *σ*_H_^2^: variance of ln(*S*_H_), (xi) *μ*_R_: mean of ln(*S*_R_), (xii) *σ*_R_^2^: variance of ln(*S*_R_).

Cell sizes at consecutive initiations are correlated as *ρ*_i_ = 1/2. Therefore the IA model can be seen as a specific case of the sHC model, for which there is no cross-correlations between physiological variables, and for which the only non-zero auto-correlation is *ρ*_i_ = 1/2. In general, *ρ*_d_ < 1/2, and it only reproduces the adder correlation in the deterministic limit where *λτ*_cyc_ is a non-stochastic variable.

### 2.3. The replication double adder (RDA) model

The RDA model states that the cell simultaneously follows two types of adder. The first adder is between two consecutive initiation cycles (‘initiation adder’), same as in the IA model. The second adder states that the size added between initiation and division is independent of the cell size at initiation (‘initiation-to-division’ adder). This second initiation-to-division adder makes the RDA model different from the IA model, although both models can be considered initiation-centric. This model was developed to explain one specific data set with non-overlapping cell cycles in *E. coli* (17). In Section 3, we will use the same statistical analysis method that was used in (17) to establish the RDA model.

In the RDA model, the cell size per origin of replication, *s*_i_, follows the following recursion relation as in the IA model (Eq. 3). As for the initiation-to-division adder, the cell size at division is determined by the following recursion relation:

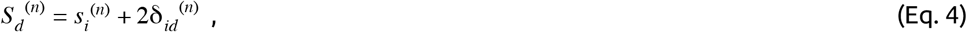

where *δ*_id_ represents the added size per origin of replication from initiation to division.

While Eq. (4) is straightforward to understand for a non-overlapping cell cycle, it is worth checking its validity for overlapping cell cycles. Let *S*_i_ ^(n)^ be the cell size at initiation for the *n*-th generation. Let us first emphasize how *S*_i_ ^(n)^ is measured. In the sHC model, *S*_i_ ^(n)^ is measured a duration of time *τ*_cyc_ ^(n)^ before division occurs. For no overlapping cell cycle, *τ*_cyc_ ^(n)^ < *τ*^*(n)*^, therefore *S*_*i*_ ^(n)^ is measured in generation *n*. For two overlapping cell cycles *τ*_*cyc*_ ^(n)^ > *τ*^*(n)*^, initiation therefore occurs in the (*n*-1)-th generation, meaning that *S*_i_ ^(n)^ refers to a size measured in the mother cell (*i*.*e*. generation *n*-1) as shown in Figure 1. Cells are born with 2 origins of replications, therefore we have *S*_i_ ^(n)^ = 2*s*_i_ ^(n)^. In this example, the mass synthesized between two consecutive replication initiation events must take into account one division event. Back to the RDA model, and using the same convention, the total added size from initiation to division is 2*S*_d_ ^(n)^ - *S*_i_ ^(n)^ = 4 *δ*_id_ ^(n)^. The factor of 4 accounts for the 4 origin of replications present after replication initiation. Dividing by 2, we obtain Eq. 4. This reasoning generalizes to any number of overlapping cell cycles. From Eq. 4, we also obtain that the average cell size at division is *S*_d_ = 2 (*δ*_ii_ + *δ*_id_). An argument similar to Eq. 2 yields the number of overlapping cell cycles *p*+1 as a function of the mean of the physiological variables: *p* is the integer part of log_2_(1 + *δ*_id_/*δ*_ii_).

The IA model is not compatible with size convergence by the adder principle. While ρ_i_ =1/2 as in the IA model, the division size mother-daughter correlation is given by

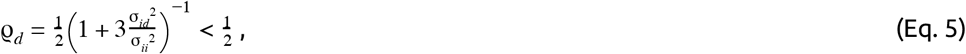

where *σ*_ii_^2^ and *σ*_id_^2^ are the variances for *δ*_ii_ and *δ*_id_ respectively (see Appendix A). Since the adder principle is equivalent to *ρ*_d_ = ½ (see Appendix A), the IA model converges to the adder only in the deterministic limit σ_id_ → 0. In addition, we can also compute the correlation between initiation size per origin and division size *ρ*_id_ = *ρ*(*s*_i_^(n)^, *S*_d_^(n)^) and obtain:

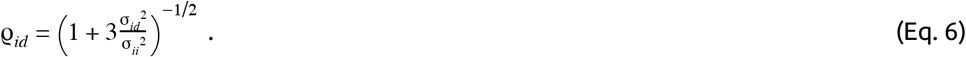

### 2.4. The independent double adder (IDA) model

The independent double adder (IDA) model states that, in steady state, initiation and division independently follow the adder principle. That is, the size added between two consecutive initiations is independent of the size at initiation (as in the IA and RDA models), whereas the size added between two division cycles is independent of the cell size at birth (or division). The recursion relation for the division size can be written as:

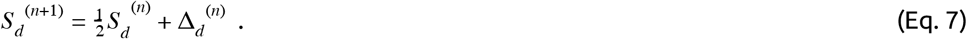

It follows that the average cell size at division is *S*_d_ = 2 *Δ*_d_. An argument similar to Eq. 2 yields the number of overlapping cell cycles *p*+1, where *p* is the integer part of log_2_(*Δ*_d_/*δ*_ii_).

We have *ρ*_i_ =1/2 and ρ_d_ = 1/2 as expected by the definition of the model. Furthermore, since initiation and division follow two independent processes (Eqs. 3 and 7), division and initiation sizes are independent from each other, namely ρ_id_ = 0.

Mechanistically, the IDA model is based on (i) balanced biosynthesis of cell-cycle proteins and (ii) their accumulation to respective threshold numbers to trigger initiation and division (11).

### 2.5. The concurrent cell-cycle processes model (CCCP)

The CCCP model is an adaptable model with several adjustable parameters (as in the sHC model) and lies somewhere in between the IA and the IDA model. The adaptability is analogous to the presentation by Amir (10) so that the model can be continuously adjusted between sizer and timer depending on the mother-daughter size correlations between -1 and +1. To ensure 1-1 correspondence between the replication cycle and the division cycle, the model explicitly implements a constraint that division must wait until after replication termination. Biologically, the model follows the view by Boye and Nordström (22).

We discuss the specific case of the adder by fixing the mother-daughter size correlation coefficients to 1/2 as explained throughout this section 2. That is, the cell size at initiation follows the recursion relation:

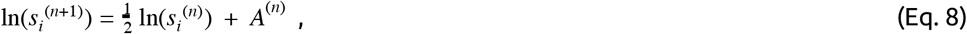

where *A*^(n)^ is the logarithmic added size between consecutive replication initiations. As mentioned above, the CCCP model was originally introduced in a more general form than Eq. 8, with an adjustable correlation parameter (see Appendix D). However, as explained by the authors a value of 1/2 is the most consistent with experimental data. Eq. 8 is very similar to Eq. 2: it is an adder on the logarithmic sizes rather than on the actual sizes at replication initiation. Denoting *C* the time to replicate the chromosome, a candidate size for the division size is:

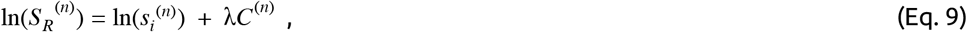

where as before *λ* is the elongation rate. If chromosome replication was the only process determining the size at division, *S*_*R*_ ^(n)^ would be the division size. However, another process, namely the division adder, is constraining the division size, resulting in a second candidate size for division:

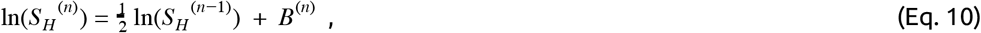

where *B* is the added logarithmic size between consecutive division adder cycles. Eq. 10 is similar to Eq. 7 and represents the division adder. Finally, cell size at division is determined by the slowest of the two processes from Eqs. 9 and 10:

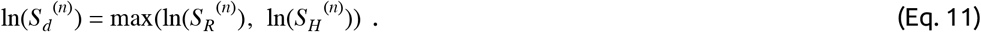

Equation 11 simply means that division should start only after replication termination. Denoting *f* as the fraction of cases in which division size is limiting (namely *S*_R_<*S*_H_=*S*_d_), the average time elapsed between replication initiation and cell division can be expressed as (assuming that < ln(*x*) >≈ ln(< *x* >) (21)):

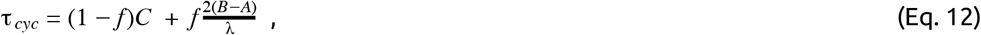

where *A, B, C* stand for means. Therefore, the number of overlapping cell cycles *p*+1 is determined by Eq. 2. Eq. 12 has a functional dependence on growth rate compatible with experimental reports (16).

### 2.6 Similarities and differences between the sHC, IA, RDA, IDA, and the CCCP models

The question of implementation point for cell size control has been controversial in the past. In the sHC, IA, and RDA models, replication initiation is the implementation point of cell size control. By contrast, the IDA and CCCP models assume that the division and replication cycles are controlled by independent processes.

These models reflect a major challenge for identifying a cell-size control model that is compatible with the new plethora of high-throughput single-cell data (2, 16). Although the sHC and IA models can be dismissed by experimental evidence (Section 3.1), the other models require more thorough analysis. For example, in contrast to the IDA model, the RDA model only ensures the initiation adder and it only reproduces the division adder behavior in the deterministic limit where *δ*_id_ is constant. The essential difference between the IDA and RDA models comes from the correlation between the size per oriC at initiation and the added size per oriC from initiation to division. Specifically, *ρ*(*s*_i_, *δ*_id_) is null for the RDA model whereas it takes negative values for the IDA.

In the next section, we test these models against data in more detail.

## 3. Test of the models against data

### 3.0 Description of the experimental data used in this study

We used datasets from our previous studies for *E. coli* and *B. subtilis* (11, 27). We have also performed additional experiments for this study (see Methods). All data and numerical analysis are available (28). In total, we have 15 experimental datasets from our studies. We have also analyzed the 4 experimental datasets made available by Witz and colleagues in their work (17).

### 3.1 Comparison of correlations vs experimental data

We first set out to test the different cell-cycle and cell-size control models. Specifically, we computed the four correlations *ρ*(*S*_b_, *Δ*_d_), *ρ*(*s*_i_, *δ*_ii_), *ρ*(*s*_i_, *δ*_id_), and *ρ*(*s*_i_, *τ*_cyc_) (Figure 2). The correlation *ρ*(*S*_b_, *Δ*_d_) is important because *ρ*(*S*_b_, *Δ*_d_)=0 defines the adder-based cell-size homeostasis. Indeed, *ρ*(*S*_b_, *Δ*_d_) is zero in virtually all experimental data. The *ρ*(*s*_i_, *δ*_ii_) is also close to zero, although deviations are seen for some experiments. These results suggest that both the IDA and RDA models are possible. By contrast, the *ρ*(*s*_i_, *τ*_cyc_) correlation shows consistently a negative value. This refutes the sHC and IA models, which both assume that the initiation-to-division duration and the initiation size per origin are independent control parameters, and thus predict *ρ*(*s*_i_, *τ*_cyc_) = 0. In addition, *ρ*(*s*_i_, *δ*_id_) is also close to zero, in favor of the RDA model, but it is also slightly negative for several conditions, in agreement with the IDA model (Appendix B). We did not test the CCCP model because it is a model to be adjusted to the data. The candidate models are therefore the RDA and IDA models. Hereafter, we therefore focus on these two models.

**Figure 2.**
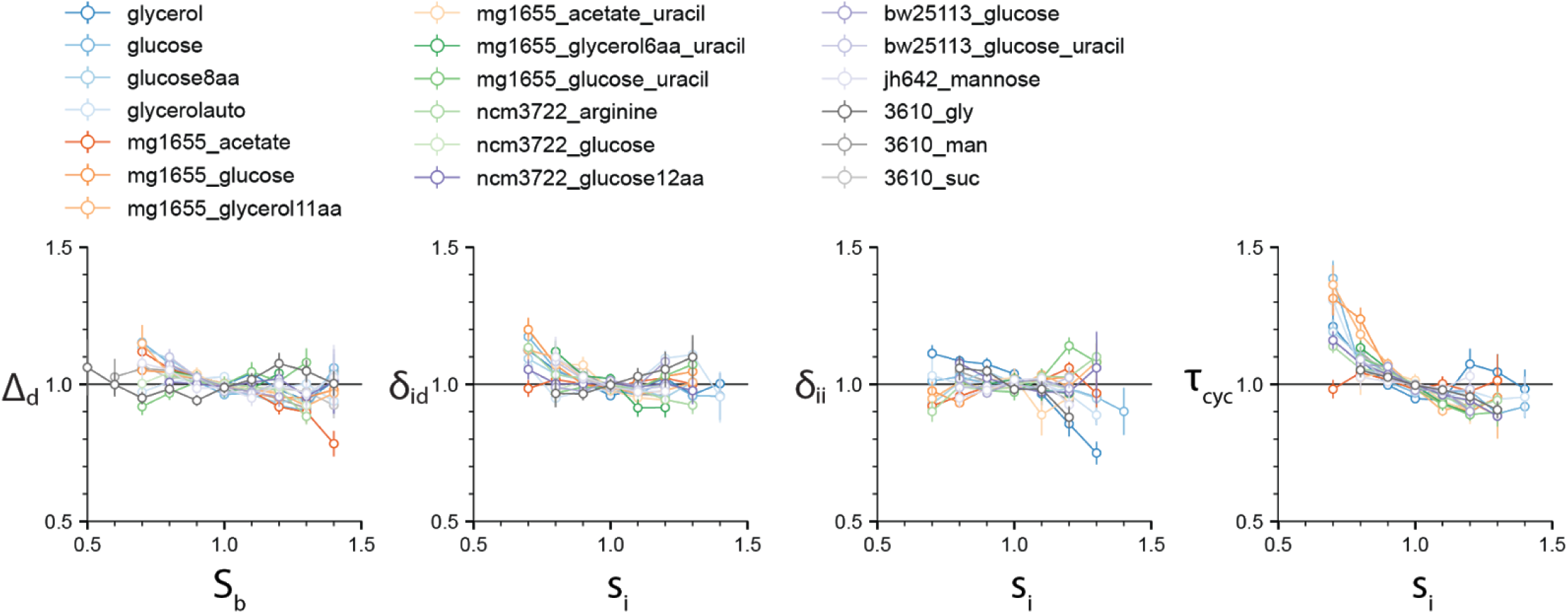
The four correlations *ρ*(*S*_b_, *Δ*_d_), *ρ*(*s*_i_, *δ*_id_), *ρ*(*s*_i_, *δ*_ii_) and *ρ*(*s*_i_, *τ*_cyc_) are computed for the 4 experimental datasets by Witz and colleagues, and 15 experimental datasets that we produced (see Methods). While the first three vanish for most experimental data, the *ρ*(*s*_i_, *τ*_cyc_) displays a consistent negative correlation, inconsistent with the HC and IA models.

### 3.2 Statistical analysis and the case study of the *I*-value analysis

In their recent paper (17), Witz *et al*. tracked replication and division cycles at the single-cell level, using experimental methods similar to previous works (11, 16, 29). They computed correlations between all pairs of measured physiological parameters, and attempted to identify the set of most mutually uncorrelated physiological variables by computing the “*I*-value”, a metric to measure the statistical independence of the measured variables. They then assumed that statistically uncorrelated physiological parameters must represent biologically independent controls. Such approaches previously facilitated the discovery of the adder principle and its formal description (8). Based on this correlation analysis or *I*-value analysis, Witz *et al*. concluded that the RDA model is the most likely model of the cell-cycle and cell-size control.

To compute the *I*-value for a given model, one needs to identify the control parameters of the model and their characteristic features. For the RDA model, they are (i) the absence of correlation between the cell size at initiation and the added size between initiation and division, namely *ρ*(*s*_i_, *δ*_id_) = 0, and (ii) the absence of correlation between the cell size at initiation and the added size between consecutive initiations, namely *ρ*(*s*_i_, *δ*_ii_) = 0. In addition to these size variables, it is known that the growth rate is mostly independent of the other physiological variables, namely: *ρ*(*λ, δ*_id_) = 0 and *ρ*(*λ, δ*_ii_) = 0. Witz *et al*. hence proposed a scalar metric that summarizes these four correlations being equal to zero, namely the determinant *I* (or *I*-value) of the matrix of correlations between the 4 variables *s*_i_, *δ*_id_, *δ*_ii_ and *λ* (Eq. 13). When *I* ≪ 1, some cross-correlations exists and both *ρ*(*λ, δ*_id_) and *ρ*(*λ, δ*_ii_) cannot vanish. On the other hand, when *I* = 1, the RDA model holds. Although since the work by Cooper and Helmstetter (23) it has been known that the progression of cell size and cell cycle can be completely described using three variables (16, 25), four variables are necessary here to encompass the correlation structure characterizing the RDA model, as explained by Witz *et al*. (17, 30). In summary, to measure the statistical independence of each set of parameters, the *I*-value analysis needs a correlation matrix of the following form (Eq. 13).

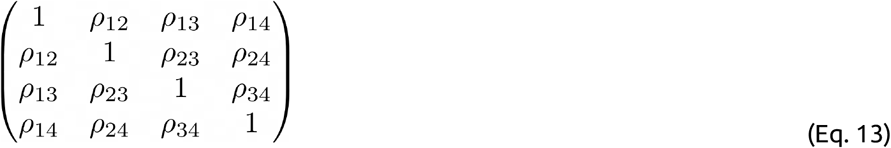

The diagonal elements are 1’s and the off-diagonal elements are cross-correlations between pairs of parameters. Therefore, if all parameters are statistically independent of each other, the off-diagonal elements should be 0, and the determinant *I* of the matrix should be 1. Based on this observation, Witz *et al*. used the determinant *I* ≤ 1 of the matrix as a metric for statistical independence of the hypothetical control parameters, with *I*=1 being the set of most independent parameters.

In Section 4, we will come back later to some of the limitations of the *I*-value analysis.

### 3.3. Test of the *I*-value analysis with various models

Our 4-variable *I*-value analysis of the models described in Section 2 (except the CCCP) for all 19 datasets is shown in Figure 3 (top). We computed *I*-values for the following four models: RDA, IDA, IA and HC (see Methods). The results of the analysis indicate that 7 out of our 15 experiments support the IDA rather than the RDA model, whereas 1 supports the IA model. Furthermore, when we applied the same analysis to all 4 datasets from Witz et *al*., we found that all 4 experiments support the IDA model. Note that Witz et al. had only analyzed one dataset (17) The sHC and IA models were included for completeness, and they show systematically a lower *I*-value (except in one *B. subtilis* condition in which the IA models had the largest score for reasons we do not understand). Overall, the results of this analysis suggest that the IDA model is most consistent with the 19 datasets we have analyzed.

**Figure 3.**
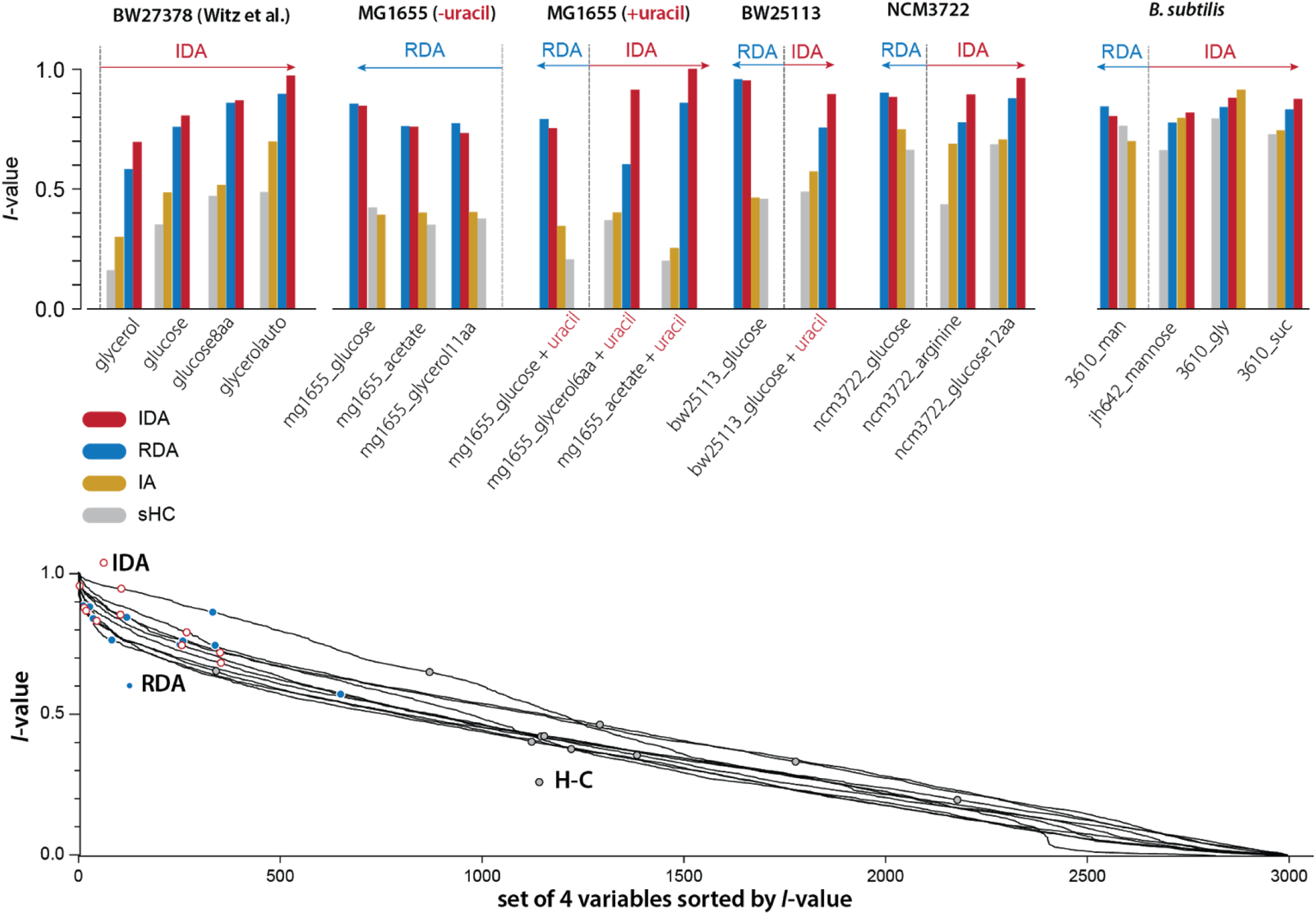
We applied the I-value analysis proposed by Witz and colleagues to 4 models (RDA, IDA, IA and HC) using 15 experimental datasets that we produced (see Methods), and the 4 datasets published by Witz and colleagues. The length of the arrows indicate how much IDA or RDA model is favored by the data.

## 4. Discussion

### 4.1 Limitations of the *I*-value analysis

We noticed that the *I*-values for the RDA and IDA models were in general very close, although these two models point to two fundamentally different mechanisms of the cell cycle. We therefore asked to what extent the *I*-value analysis could be used to effectively identify meaningful models of the cell cycle. In the spirit of the ranking performed in the study by Witz et *al*., we considered all possible combinations of 4 among 18 physiological variables (see Methods; see Figure S1), and computed the *I*-values for each of Witz and colleagues’ 4 datasets and for each of our 15 datasets. Although the IDA and RDA models have high scores, we found that many other combinations have higher *I*-values, including combinations that do not correspond to any meaningful model of the cell cycle (Figure 3 bottom). Since *I*-values cannot be used to distinguish sound from unsound models of the cell-cycle, we conclude that this analysis lacks predictive power.

Furthermore, the *I*-value analysis can only be employed to compare models with the same number of variables in defining correlations. To see this, let us consider the RDA and IDA models in Figure 4. The RDA model can be defined by the 3 parameters {*δ*_ii_, *δ*_id_, *λ*} (Figure 4a). Indeed, from an initial condition consisting of an initiation size, only those 3 parameters need to be known at each generation to construct a whole lineage. Yet the defining correlations of the RDA model are given by *ρ*(*λ, δ*_id_) = 0 and *ρ*(*λ, δ*_ii_) = 0. Thus, we need a total of 4 variables {*s*_i_, *δ*_ii_, *δ*_id_, *λ*} to characterize the RDA model, which leads to the 4×4 correlation matrix shown in Figure 4c.

**Figure 4.**
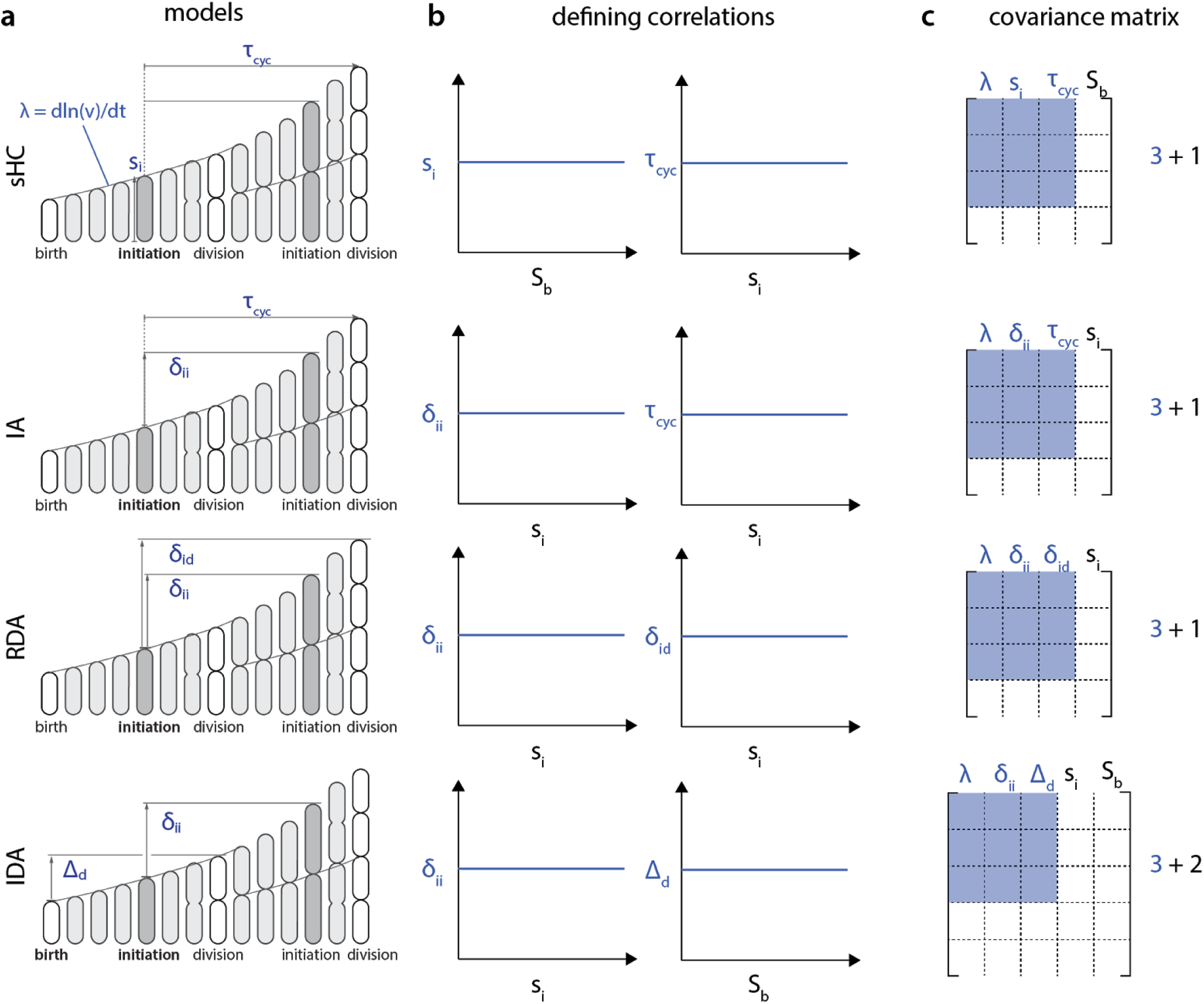
Each model (**a**) is characterized by a set of two correlations (**b**). Note that these defining correlations require 1 additional parameter *S*_b_ for the sHC model, 1 additional parameter *s*_i_ for the RDA and the IA models, whereas the IDA model requires 2 additional parameters, *s*_i_ and *S*_b_. **c**. As a result, the covariance matrix for the *I*-value analysis of these models, according to Witz *et al*., would be 4×4 for the sHC, IA and RDA models and 5×5 for the IDA model. Therefore, the *I*-values of these models cannot be meaningfully compared.

The problem with the above procedure is that the size of the correlation matrix becomes model dependent, and thus the *I*-value analysis cannot compare different models. For example, following the same reasoning, the IDA model would require 5 variables for the *I*-value analysis, because of the two defining correlations (*s*_i_, *δ*_ii_) and (*S*_b_, *Δ*_d_) (Figure 4b). Therefore, in addition to the three independent control parameters {*δ*_ii_, *Δ*_d_, *λ*}, the *I*-value analysis would require two additional parameters {*s*_i_, *S*_b_} from the defining correlations. The resulting correlation matrices would then be 5×5 from {*s*_i_, *S*_b_, *δ*_ii_, *Δ*_d_, *λ*} instead of 4×4 (Figure 4c). Since *I*-values obtained from correlation matrices of different sizes cannot be meaningfully compared, the *I*-value analysis is fundamentally limited to compare a specific class of models.

Finally, it is worthwhile mentioning that the *I*-value analysis employed by Witz *et al*. is only valid for non-overlapping cell cycles (30).

### 4.2 The RDA model does not produce the adder principle

From Eq. 5, size convergence according to the RDA model is incompatible with the adder principle. More specifically, in the presence of fluctuations, the RDA model is skewed toward a sizer behavior. Using experimentally measured values for the variance of *δ*_ii_ and *δ*_id_, we computed *ρ*_d_ according to Eq. 5 (Figure 5). The RDA model would predict a deviation from the adder principle, in contradiction to several experimental results (6, 8, 31, 32)

**Figure 5:**
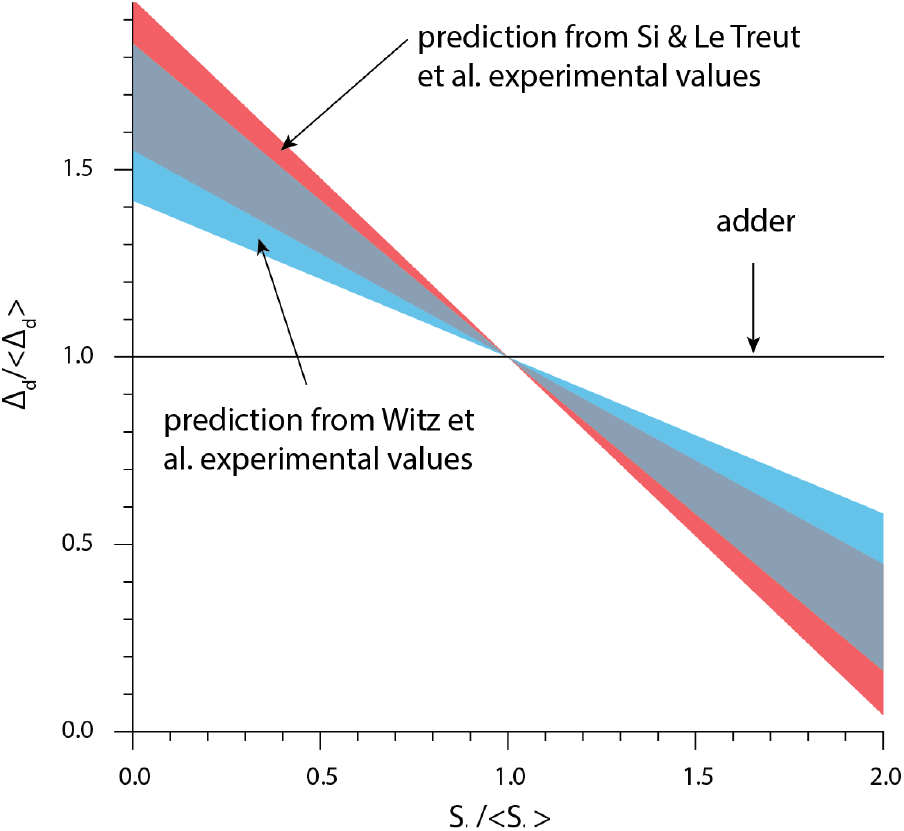
The theory based on the initiation-centric model predicts a more sizer-like behavior. We used Eq. 1 and the experimental values of σ_id_/σ_ii_ from (17) and (11).

Witz et *al*.’s simulation of the RDA showed a good agreement with the experimental data (17). Yet further investigation showed that the agreement was a direct consequence of introducing yet another adjustable parameter, namely the variance of the septum position (31, 32). Indeed, for perfectly symmetric division, the simulation results also show deviatiations from experimental adder behavior (Figure S2), in agreement with Eq. 5.

### 4.3 Mechanistic origin of the IDA model

The IDA model is a mechanistic model based on two experimentally verified hypotheses. First, the cell cycle proteins are produced in a balanced manner (i.e., the synthesis rate of each proteins is the same as the growth rate of the cell):

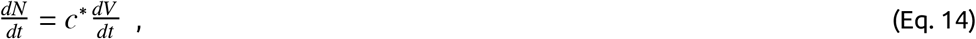

where *N* is the protein copy number in the cell, *V* is the cell volume and *c*\* is the steady-state protein concentration. Second, initiation or division is triggered when the respective initiator protein reaches a threshold, namely:

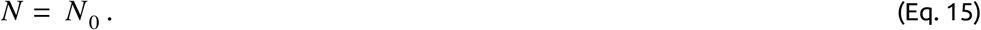

The adder phenotype is a natural consequence of these two assumptions, provided that the initiator proteins are equally partitioned at division between daughter cells (11). The requirement that *N*_*0*_*/2* proteins must be synthesized between birth and division results in the added volume from birth to division to be:

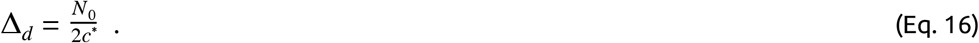

This model was substantiated in an experimental study showing that perturbing the first condition, namely balanced biosynthesis, was enough to break the adder phenotype (11). Balanced biosynthesis was perturbed in two orthogonal ways: (i) by oscillating the production rate of the FtsZ protein through periodical induction and (ii) by relieving FtsZ degradation through ClpX inhibition.

A similar mechanism is thought to apply to the initiation process, through the initiator protein DnaA, which accumulates at the origin of replication to trigger replication initiation. An important difference with the division mechanism however is that this results in a threshold to be reached at each origin of replication, thus (Eq. 14) is modified to:

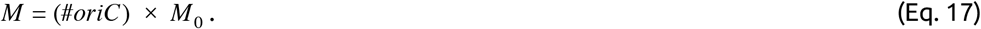

Provided that once again proteins are equally partitioned at division, Equations (2) and (4) result in a fixed added volume per origin of replication between consecutive initiation events, hence the initiation-to-initiation adder *δ*_ii_.

#### Perspective and concluding remarks

While applying a recently proposed correlation analysis, namely the *I*-value analysis, we became aware of some of its limitations, as explained above. In our view, this illustrates some of the caveats one may encounter when applying correlation analysis. While valuable in various contexts, correlation analysis can lead to erroneous conclusions when additional sources of variability such as experimental and measurement errors are not properly taken into account in the analysis. For example, adder correlations can emerge from non-adder mechanisms due to measurement errors in the cell radius (33). Therefore, while correlation analysis is useful to confront models to experimental data, we believe it is important to seek a molecular understanding of a model of the cell cycle.

The question of whether the implementation point of the cell cycle is birth or replication initiation has a long history. Although cell-size control was initially thought to be division centric because the CV of the division size was smaller than that of the doubling time (34), many interpreted the HC model (23) and Donachie’s theoretical observation (24) as an initiation-centric view for long. The rediscovery of the adder principle, not explained within the HC model, has revealed the need to revisit models of the bacterial cell cycle. In this article, we have reviewed one of the latest controversies that has emerged in the field of quantitative bacterial physiology, namely the question of the implementation point of the cell cycle. Based on recent results and single-cell experimental data that we have generated over the last decade, we favor a mixed implementation strategy with two independent adders namely the IDA, as the most likely mechanism ruling the *E. coli* and *B. subtilis* cell cycle. Furthemore, the fact that both initiation and division share the same adder phenotype suggest to us that they also must share the same mechanistic principles. That is, initiation and division must require (i) balanced biosynthesis of their initiator proteins such as DnaA for initiation and FtsZ for division, and (ii) their accumulation to a respective threshold number to trigger initiation and division (11). Ultimately, these predictions should be tested experimentally to gain mechanistic understanding and their generality beyond correlation analysis.

## Materials and Methods

### Strains and growth conditions

The following strains were used in this study.

- BW25113: F-DE(araD-araB)567 lacZ4787(del)::rrnB-3 LAM-rph-1 DE(rhaD-rhaB)568 hsdR514.

(Note that the genotype of BW27378 is F-DE(araD-araB)567 lacZ4787(del)::rrnB-3 LAM-rph-1 DE(rhaD-rhaB)568 hsdR514 DE(araH-araF)570(::FRT).)

- SJ_FS130: To construct this strain, we introduced ΔdnaN::[dnaN-ypet] into BW25113 using P1 transduction.
- SJ_DL188: MG1655 F-λ-rph-1 dnaA msfGFP kan mCherry-dnaN

We used minimal MOPS media for the MG1655 experiments and minimal M9 glucose media with and without uracil for the SJ_FS130 (BW25113 based) experiments.

### Microfluidics, Microscopy and Image processing

We used the same method as described in (11).

## Appendix A: Cell size homeostasis in the RDA model

In the model proposed by Witz and colleagues, the cell size per origin at division is determined by the cell size at initiation per origin *s*_i_, the added size per origin between consecutive replication initiation events *δ*_ii_, and the added size per origin from replication initiation to cell division *δ*_id_. The following relation holds:

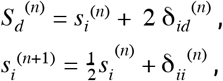

where the index as *n* denotes the generation (or division cycle) . Under the assumption that *δ*_ii_ ^(n)^ (resp. *δ*_id_ ^(n)^) are independently and identically distributed Gaussian stochastic variables with mean *μ*_ii_ (resp. *μ*_id_) and standard deviation *σ*_ii_ (resp. *σ*_id_), it follows that *s*_i_ ^(n)^ and *S*_d_ ^(n)^ are also Gaussian stochastic variables. At large *n*, they converge to the limiting distributions *s*_i_ ≡ N(*μ*_i_, *σ*_i_) and *S*_d_ ≡ N(*μ*_d_, *σ*_d_), where:

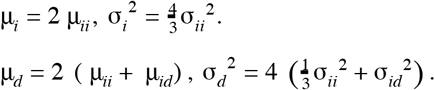

The mother-daughter correlation for division size is a central quantity in cell size homeostasis, which can be derived in this model. As a first step, let us define the centered variables: d*s*_i_ ^(n)^ = *s*_i_ ^(n)^ - μ_i_ and d*S*_d_ ^(n)^ = *S*_d_ ^(n)^ - *μ*_d_ . We then obtain the relations:

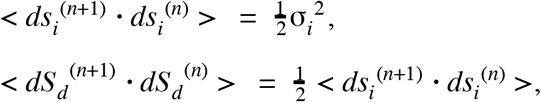

where the brackets denote averages. We therefore obtain the mother-daughter Pearson correlation coefficients (in the large *n* limit):

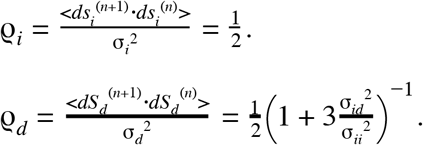

It also follows that the Pearson correlation coefficient between initiation size per origin and division size is:

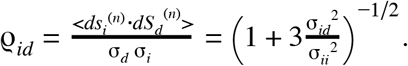

In this model the joint distribution (*S*_d_ ^(n)^, *S*_d_ ^(n-1)^) is a bivariate Gaussian, therefore we can write the conditional expectation of *S*_d_ ^(n)^ as:

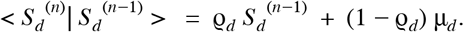

With the hypothesis of symmetrical division, namely S_b_ ^(n)^ = 2 S_d_ ^(n-1)^, we obtain for the conditional expectation of the added size from birth to division:

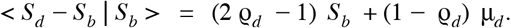

Therefore, the “adder” principle is equivalent to having *ρ*_d_ = 1/2. Therefore, the RDA model always results in 2 *ρ*_d_ - 1 < 0. In fact, it only reproduces the “adder” principle in the deterministic limit σ_id_ → 0.

## Appendix B: Calculation of *ρ*(*s*_i_, *δ*_id_) for the IDA model

In the IDA model, the cell size at division is determined by the cell size at birth *S*_b_ and the added size from birth to division *Δ*_d_; and the cell size per origin at initiation in the next cell cycle *s*_i_ ^(n+1)^ is determined by the added size per origin between consecutive replication initiation events *δ*_ii_, and the cell size per origin at initiation *s*_i_. The following relations hold:

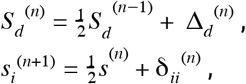

where the index *n* denotes the generation (or division cycle), and {Δ_d_ ^(n)^} and {*δ*_ii_ ^(n)^} are independently distributed random variables. Denoting μ_*dd*_ = < Δ_*d*_ > and μ_*ii*_ = < δ_*ii*_ >, we have μ_*d*_ = < *S*_*d*_ > = 2 μ_*dd*_ and μ_*i*_ = < *s*_*i*_ > = 2 μ_*ii*_ . We also define the centered variables: *d*Δ _*d*_^(*n*)^ = Δ_*d*_ ^(*n*)^ − μ_*dd*_, *dS*_*d*_ ^(*n*)^ = *S*_*d*_ ^(*n*)^ − μ_*d*_, *d*δ_*ii*_ ^(*n*)^= δ_*ii*_^(*n*)^ −μ_*ii*_ and *ds*_*i*_^(*n*)^ = *s*_*i*_^(*n*)^ −μ_*i*_ . Denoting σ_*dd*_ ^2^ = < *d*Δ_*d*_ ^2^ > and σ_*ii*_ ^2^ = < *d*δ_*ii*_ ^2^ >, we have 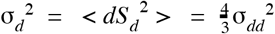 and 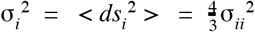.

The added size between initiation and division reads δ_*id*_ ^(*n*)^ = (*S*_*d*_ ^(*n*)^ −*s*_*i*_ ^(*n*)^)/2, where the factor of 2 reflects the fact that origins of replication double at initiation. Therefore:

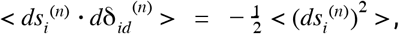

by independence of *S*_d_ ^(n)^ and *s*_i_ ^(n)^, and we obtain the Pearson correlation coefficient:

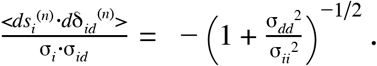

## Appendix C: Calculation of *ρ*(*s*_i_, *S*_d_) and *ρ*(*S*_d_ ^(n)^, *S*_d_ ^(n+1)^) for the sHC and IA models

Let us rewrite Eq. 1 as:

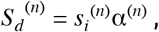

Where α^(*n*)^ = exp(λ^(*n*)^τ _*cyc*_^(*n*)^) . To derive *ρ*(*s*_i_, *S*_d_) and *ρ*(*S*_d_ ^(n)^, *S*_d_ ^(n+1)^), we will assume that {*s*_i_ ^(n)^} and {*α*^(n)^} are independent random vectors. This is true in the IA model (because *δ*_ii_ is independent of the other physiological variables), but it is an additional constraint for the sHC model. Using this assumption, we can derive the mean and variance of the cell size at division. The mean is given by:

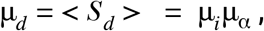

where μ_*i*_ =< *s*_*i*_ > and μ_α_ =< α > . Let us introduce the centered variables: *ds*_*i*_^(*n*)^ = *s*_*i*_^(*n*)^ −μ_*i*_, *d*α^(*n*)^ = α^(*n*)^ −μ_α_ and *dS*_*d*_ ^(*n*)^ = *S*_*d*_ ^(*n*)^ −μ_*d*_, and the variances σ_*i*_ ^2^ =< *ds*_*i*_ ^2^ > and σ_α_^2^ =< *d*α^2^ > . The variance of the cell size at division is given by:

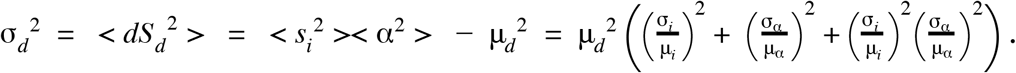

We now compute *ρ*(*s*_i_, *S*_d_):

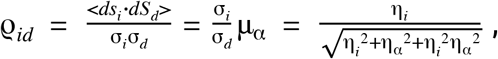

where we have introduced the coefficient of variations (CVs): η_*i*_ = σ_*i*_/μ_*i*_ and η_α_ = σ_α_/μ_α_ . As can be seen, the value of *ρ*_id_ depends on the CVs of the initiation size and of α = exp(λτ _*cyc*_) . When *α* is deterministic (*i*.*e. η*_*α*_=0), division is “slaved” to replication initiation and *ρ*_id_=1.

To compute *ρ*(*S*_d_ ^(n)^, *S*_d_ ^(n+1)^), we introduce the mother/daughter correlations:

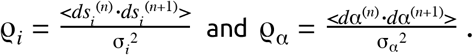

In particular, we have < *s*_*i*_^(*n*)^ ° *s*_*i*_^(*n*+1)^ > = μ_*i*_^2^ + ϱ_*i*_σ_*i*_^2^ and < α^(*n*)^ ° α^(*n*+1)^ > = μ_α_^2^ + ϱ_α_σ_α_^2^ . We thus have:

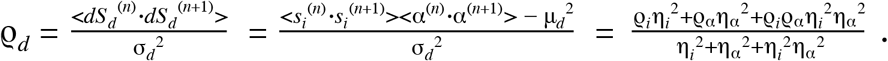

When *α* is deterministic (*i*.*e. η*_*α*_=0), we have *ρ*_d_ = *ρ*_i_: cell size homeostasis is determined by the initiation size homeostasis. On the other hand, when *s*_i_ is deterministic (*i*.*e. η*_*i*_=0), we have *ρ*_d_ = *ρ*_*α*_. In general, the level sets of *ρ*_d_ are determined by a quadratic equation of the variables *ρ*_i_ and *ρ*_*α*_. Therefore each level set is a conic section.

In the absence of mother/daughter correlations, namely *ρ*_i_ = *ρ*_*α*_ = 0, then *ρ*_d_ = 0, which is a sizer regime. If we only have *ρ*_i_ = 0, then:

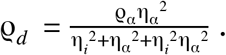

For the IA model, we have *ρ*_i_ = 1/2. Furthermore, there are no mother/daughter correlations among the other physiological variables, therefore *ρ*_*α*_ = 0. We thus obtain:

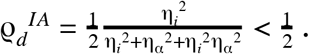

Therefore, the IA model can only reproduce the adder correlation in the deterministic limit where α = exp(λτ _*cyc*_) is a deterministic variable, namely *η*_α_=0.

## Appendix D: Calculation of *ρ*(*s*_i_, *S*_d_) and *ρ*(*S*_d_ ^(n)^, *S*_d_ ^(n+1)^) for the CCCP model

For simplicity, and following the conventions used by the authors of this model (14), we rewrite Eq. (8-11) as:

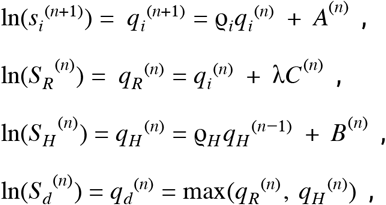

where we have introduced *ρ*_i_ and *ρ*_H_ to be more general. Assuming that *A*^(n)^, *B*^(n)^ and *C*^(n)^ are Gaussian variables with means *μ*_A_, *μ*_B_ and *μ*_C_, and variances *σ*_A_ ^2^, *σ*_B_ ^2^ and *σ*_C_ ^2^, then *q*_i_ ^(n)^, *q*_R_ ^(n)^ and *q*_H_ ^(n)^ are also Gaussian variables. We first compute their means and variances. We have:

- *q*_*i*_ ≡ *N* (μ_*i*_, σ_*i*_) with 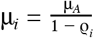 and 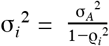;
- *q*_*R*_ ≡ *N* (μ_*R*_, σ_*R*_) with μ_*R*_ = μ_*i*_ + λμ_*c*_ and σ_*R*_ ^2^ = σ_*i*_^2^+ λ^2^ σ_*c*_^2^;
- *q*_*H*_ ≡ *N* (μ_*H*_, σ_*H*_) with 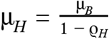 and 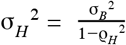.

Note that *q*_H_ is independent from *q*_i_ and *q*_R_. We now compute the autocovariances for these 3 variables. We obtain:

- < *dq*_*i*_^(*n*+1)^ *dq*_*i*_^(*n*)^ > =ϱ_*i*_ σ_*i*_^2^;
- < *dq*_*R*_^(*n*+1)^ *dq*_*R*_^(*n*)^ > = ϱ σ_*i*_ ^2^ since *C*^(n)^ is an independently distributed random variable;
- < *dq*_*H*_^(*n*+1)^ *dq*_*H*_^(*n*)^ > = ϱ_*H*_ σ_*H*_ ^2^,

where as before *dX* ^(*n*)^ = *X*^(*n*)^ −< *X* > . We furthermore compute for subsequent use the following cross-correlation: < *dq*_*R*_^(*n*)^*dq*_*i*_^(*n*)^ > = σ_*i*_^2^ since *C*^(n)^ is an independently distributed random variable. These preliminary results will be useful to compute *ρ*_d_, *ρ*_i_ and *ρ*_id_.

We now follow the approach of the authors, and replace the expression of *q*_d_ in Eq. (11) by the effective equation:

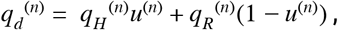

where {*u*^(n)^} are independent Bernoulli variables such that: *Pr*(*u*^(*n*)^ = 1) = *f* . We note that *Pr*(*u*^(*n*)^ = 0) = 1 − *f* and *Pr*((*u*^(*n*)^)^2^ = 1) = *f* . The previous equation is an effective approach in which *f* represents the fraction of the cases in which the division process is limiting, namely *q*_*R*_^(*n*)^ < *q*_*H*_ ^(*n*)^ . With this effective expression, one can compute the mean *μ*_d_ and the variance *σ*_d_ ^2^ of *q*_d_ . We obtain:

- μ_*d*_ = μ_*H*_ *f* + μ_*R*_(1 − *f*);
- σ_*d*_ ^*2*^=< (*q*_*H*_ ^*(n)*^)^2^ >< (*u*^*(n)*^)^2^ > + < (*q*_*R*_^*(n)*^)^2^ >< (1 −*u*^(*n*)^)^2^> + 2 < *q*_*H*_^(*n*)^ >< *q*_*R*_^(*n*)^ >< *u*^(*n*)^ (1 −*u*^*(n)*^) >− μ_*d*_ ^2^, = σ_*H*_ ^2^*f* + σ_*R*_ ^2^(1− *f*) + *f* (1− *f*)(μ_*H*_ −μ_*R*_)^2^.

We now turn our attention to the computation of *ρ*_id_. For that purpose, we compute the average:

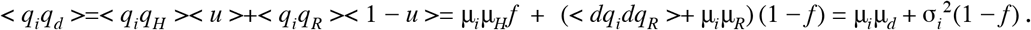

We thus obtain:

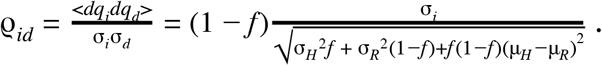

We now turn our attention to the computation of *ρ*_d_. We start by computing the average:

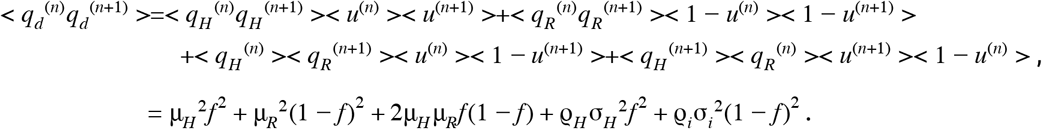

We therefore obtain:

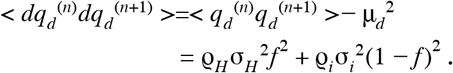

Finally, we obtain:

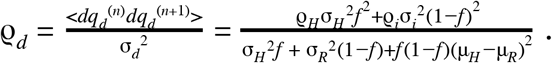

When the division process is limiting, namely *f=1*, then we have *ρ*_d_=*ρ*_H_. Conversely when the initiation process is limiting, we have *ρ*_d_=*ρ*_i_. More generally, the level set curves for *ρ*_d_ are lines in the (*ρ*_H_,*ρ*_i_) plane.

## Appendix E: *I*-value analysis

Following the methodology proposed by Witz *et al*. (17), *we first defined the covariance matrix (Eq. 13) of the analyzed variables. Then the I*-value was computed as:

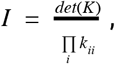

where the *k*_ii_ are the diagonal elements of the covariance matrix, hence the variances. To compute the *I*-values of the four models listed in Figure 3 (top), we used the following combinations of physiological variables:

| Model | Variables |
| --- | --- |
| RDA | $\lambda, \delta_{id}, \delta_{ii}, s_i$ |
| IDA | $\lambda, \Delta_d, \delta_{ii}, S_b$ |
| IA | $\lambda, \delta_{ii}, \tau_{cyc}, s_i$ |
| sHC | $\lambda, \tau_{cyc}, s_i, S_b$ |

In Figure 3(bottom), we defined 18 physiological variables in line with the definitions given by (17)(see Figure S1): *λ, S*_b_, *S*_d_, *S*_i_, *Δ*_d_, *Δ*_bi_, *s*_i_, *s*_i_ ^(n-1)^, *s*_i_ ^(n+1)^, *δ*_ii_, *δ*_id_, *R*_ii_, *R*_id_, *R*_bd_, *R*_bi_, *τ, τ*_cyc_ and *τ*_ii_. We generated all possible combinations of 4 variables, that is = 3060. We emphasize the difference between *s*_i_ and *S*_i_. The former is the cell size per origin of replication at the initiation event associated with the division in the current generation; therefore it could be in a previous generation (*e*.*g*. mother, grand-mother cell). The latter is the cell size when replication initiation occurs in the current generation. These 2 variables are only the same in the case of non-overlapping cell cycles (Figure 1). We also found that the results of this analysis were sensitive to processing of the experimental data. In their work, Witz and colleagues, instead of using the measured values for *S*_b_, *S*_d_, *S*_i_ and *s*_i_, first fitted traces of cell sizes to an exponential function and took values interpolated by this fit. Yet it did not affect the relative ranking of the division-centric model with respect to the initiation-centric model.

**Figure S1:**
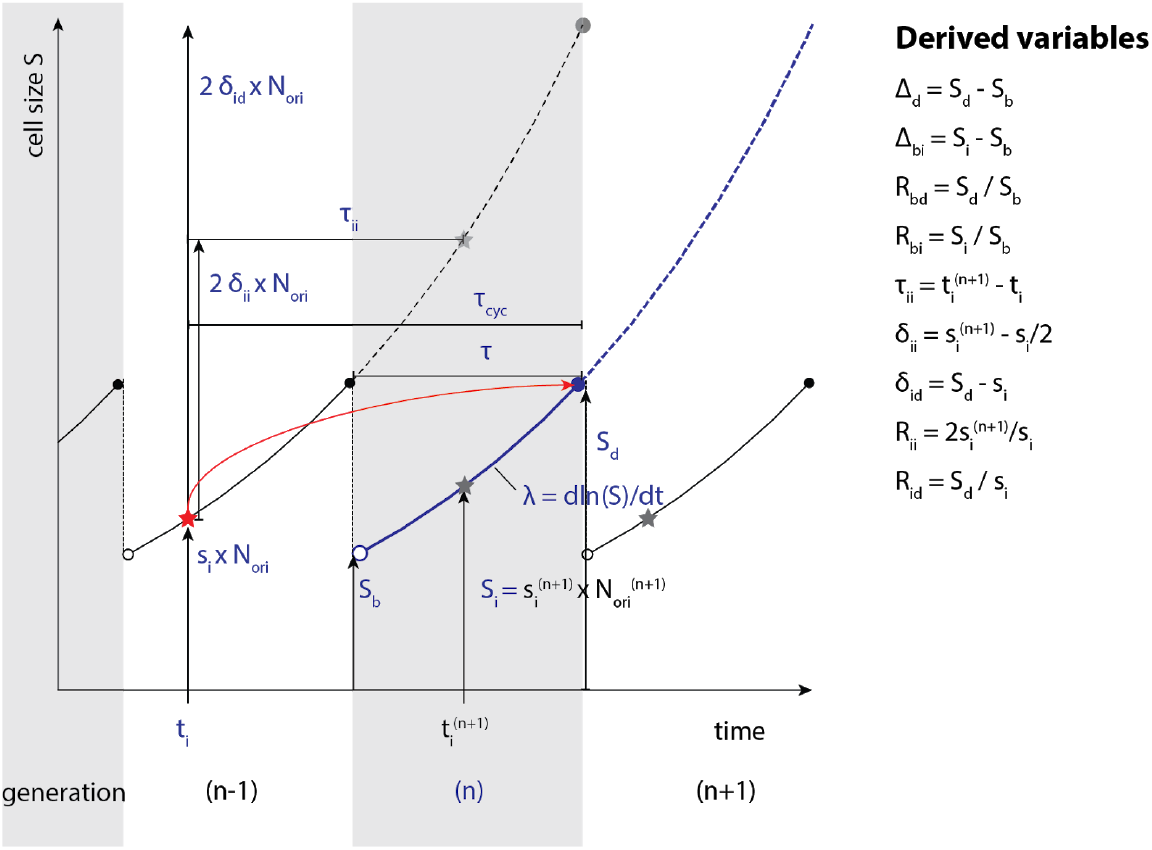
Definition of the physiological variables in a scenario with 2 overlapping cell cycles. All variables with blue font are associated with the current generation (*n*). We added a superscript when using physiological variables associated with another generation. We also defined variables derived from these quantities. Replication initiations are indicated with stars, and the red star is the initiation determining the division for the current generation. *N*_ori_ is the number of origins of replication just before initiation happens.

## Appendix F: Simulations

In this study we have performed simulations of both initiation-centric and division-centric models. For this, we have re-used the code provided in (17). Few and minor modifications have been made, but these modifications did not affect the outcome or the essence of the original simulations. Simulations performed consist of:

- Repeats of the simulations performed by Witz and colleagues in their original study.
- Simulations using Witz and colleagues’ original parameters, but with perfectly symmetrical division_i_.
- Simulations of Witz et al. model, and our model, using experimental parameters taken from experimental datasets from Witz and et al. and datasets published in (11).

Witz et al.’s simulations of the RDA reproduced the adder behavior observed in their data, in apparent contradiction with our prediction in Eq. 5 that the RDA model is inconsistent with size homeostasis by the adder. We analyzed their simulations and found that they produced the adder-like behavior because of the additional fluctuations in the septum position (Figure S2a). Experimentally, septum position represents the most precise control among all measured single-cell parameters with CV < 5% (8, 27). Indeed, removing fluctuations in the septum position alone made the simulation deviate from experiment in a quantitative manner consistent with Eq. 5. We also conducted a similar analysis using our experimental data (11), and reached the same conclusion (Figure S2b). Based on this observation, we conclude that the RDA model does not self-consistently explain the adder phenotype.

**Figure S2:**
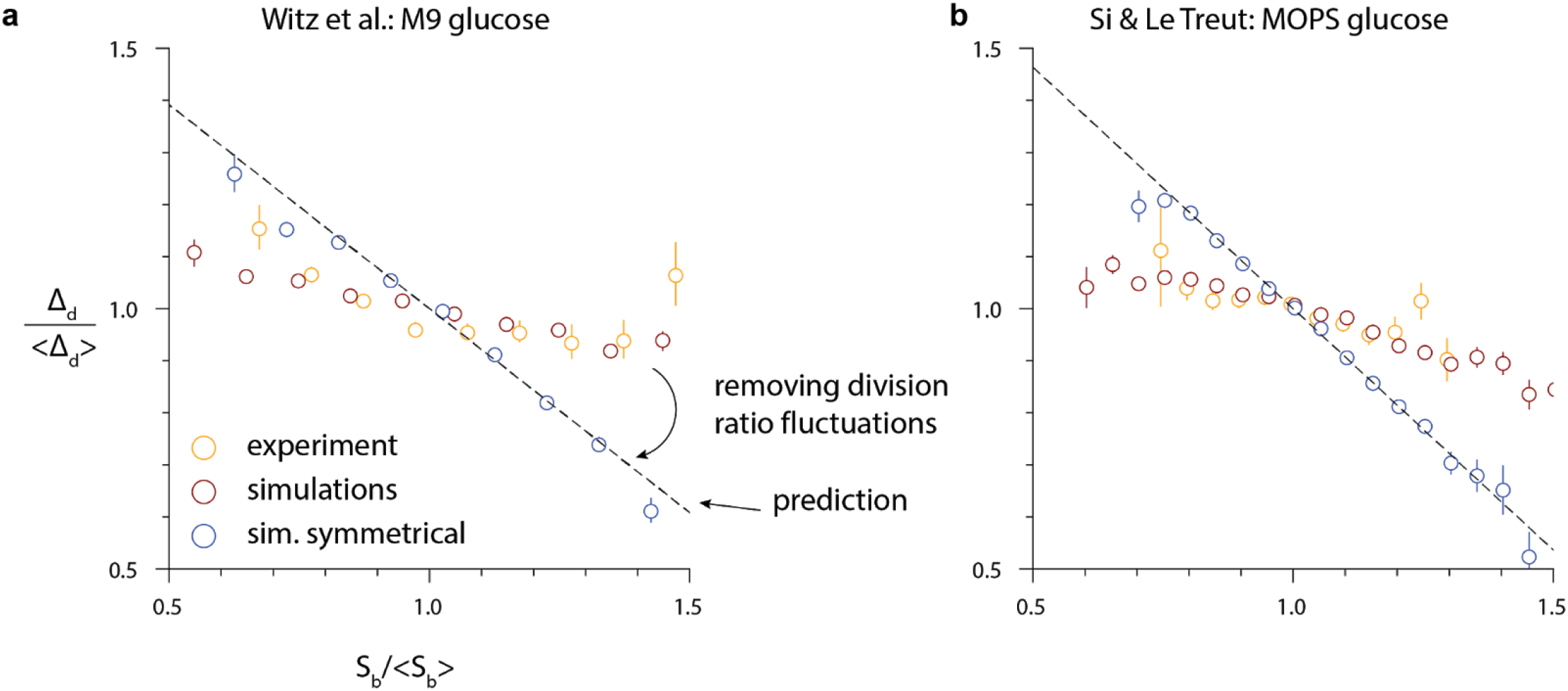
Agreement of the RDA model with experimental data from **a**. (17), M9 + glucose condition and **b**. (11), MOPS + glucose condition (MG1655 *E. coli*). After removing fluctuations in the division ratio, simulations don’t agree with experimental data.

## Supplementary Figures

**Figure S3.**
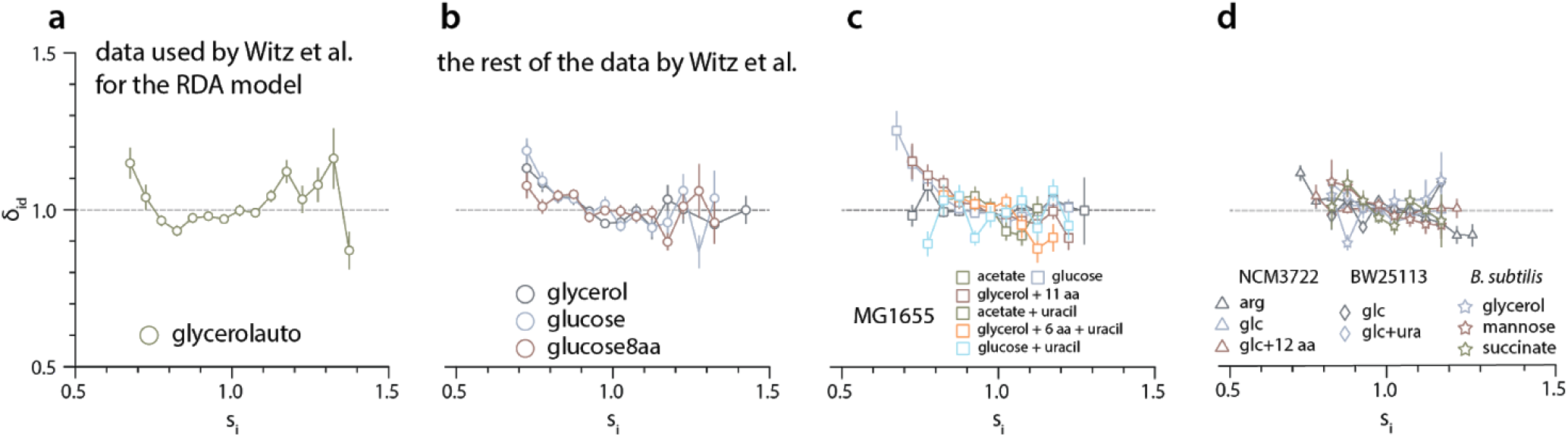
**a**. Witz et al.’s validation of the RDA model is exclusively based on this data set. **b**. 3 out of the 4 datasets of Witz et al. present a slight negative correlation. **c-d**.Most of the experimental data we obtained with *E. coli* MG1655, NCM3722, BW25113 and *B. subtilis* NCIB 3610 strain also show varying degrees of negative correlation, consistent with the IDA model.

**Figure S4.**
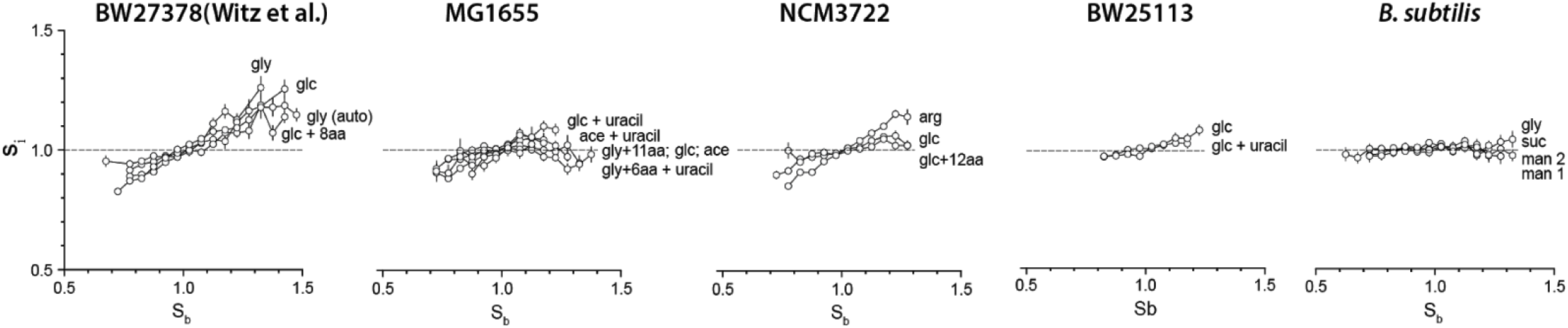
Varying degrees of positive correlation between the birth size *S*_b_ and the initiation size per origin *s*_i_. The experimental data from (17) shows the most positive correlation whereas experimental data acquired in our lab with *E. coli* MG1655 and with *B. subtilis* show moderate to zero correlation.

## Notes

### Competing Interest Statement

The authors have declared no competing interest.

